# Transcriptome Profiling of different types of human respiratory tract cells infected by SARS-CoV-2 Highlight an unique Role for Inflammatory and Interferon Response

**DOI:** 10.1101/2020.11.15.383927

**Authors:** Minghui Yang, Luping Lei, Qiumei Cao, Yang Yang, Jun Wang, Xiao Jiang, Kun Huang, Jinzhi Lai, Ling Qing, Yu Wang, Yingxia Liu

**Author notes:** Contributed equally (Minghui Yang, Luping Lei). Contributed equally (Yu Wang, Yingxia Liu). Corresponding Author: Yingxia Liu.

## Abstract

The emergence of severe acute respiratory syndrome coronavirus 2 (SARS-CoV-2) disease (COVID-19) at the end of 2019 has caused a large global outbreak and now become a major public health issue. Lack of data underlying how the human host interacts with SARS-CoV-2 virus. In the current study, We performed Venn-analysis, Gene ontology (GO), KEGG pathway analysis and Protein-protein interaction analysis of whole transcriptome studies with the aim of clarifying the genes and pathways potentially altered during human respiratory tract cells infected with SARS-CoV-2. We selected four studies through a systematic search of the Gene Expression Omnibus (GEO) database or published article about SARS-CoV-2 infection in different types of respiratory tract cells. We found 36 overlapping upregulated genes among different types of cells after viral infection. Further functional enrichment analysis revealed these DEGs are most likely involved in biological processes related to inflammatory response and response to cytokine, cell component related to extracellular space and I-kappaB/NF-kappaB complex, molecular function related to protein binding and cytokine activity. KEGG pathways analysis highlighted altered conical and casual pathways related to TNF, NF-kappa B, Cytokine-cytokine receptor interaction and IL17 signaling pathways during SARS-CoV-2 infection with CXCL1, CXCL2, CXCL3, CXCL8, CXCL10, IL32, CX3CL1, CCL20, IRF1, NFKB2 and NFKB1A up-regulated which may explain the inflammatory cytokine storms associated with severe cases of COVID-19.

## INTRODUCTION

The outbreak of Coronavirus Disease 2019 (COVID-19) caused by the novel coronavirus named severe acute respiratory syndrome coronavirus-2 (SARS-CoV-2), was first reported at the end of 2019 [1,2]. The clinical characteristics of COVID-19 include respiratory symptoms, fever, cough, dyspnea, pneumonia [3-6], acute respiratory distress syndrome (ARDS), and even multiple organ dysfunction in severe cases [7]. Clinical characteristic varied in different COVID-19 patients, and a number of patients showed asymptomatic infection [8-10] Recent studies have revealed that asymptomatic cases could also transmit the virus to contacts around them, which makes it more difficult to control the spread of the virus [11].

Several anti-viral drugs have been used to target the SARS-CoV-2 infection. These drugs include lopinavir/ritonavir, penciclovir, chloroquine, hydroxychloroquine, remdesivir, ribavirin [12]. However, most of drugs have limited efficacy to reduce the severity of COVID-19 patients. In addition, some of them have serious side effects [13,14]. The shortcomings of current tested drugs lead scientists and physicians continuously looking up the better remedies for treatment of COVID-19. Transcriptional profiling has provided remarkable opportunities for understanding the relationship between cellular function and metabolic pathways, as well as to define the possible implications of genetic variability and environmental conditions in many tissues and organisms after viral infection [15,16], which could contribute to the screening of drug targets against viruses. In bioinformatics, RNA-Seq analyses are challenging due to its complexity and voluminosity. However, many computational methods and techniques have been developed for the use of the analysis of transcriptomics data. In addition, network-based approach is also extensively used to analyze and identify the potential candidate genes after viral infection [17].

The clinical picture of COVID-19 patients are characterized by an over-exuberant immune response with lung lymphomononuclear cells infilteration and proliferation that may account for tissue damage more than the direct effect of viral replication [18-20]. Evidence also showed that SARS-CoV-2 infects human respiratory tract cells, prompting inflammation and tissue damage/loss as well as in other different cell types [21]. Despite recent advances, little is known about the pathways involved in viral pathogenesis [22]. In the present study, we performed an analysis of several transcriptome studies, aiming to clarify which genes and cellular networks were up- or down-regulated during SARS-CoV-2 infection in human lung cells. Moreover, we assessed a comprehensive GO enrichment and KEGG pathway analysis to predict how the modulation of these genes could affect the outcome of the disease.

## MATERIALS AND METHODS

### Study Search and RNA-Seq data collection

We searched for RNA-Seq experiments deposited in the Gene Expression Omnibus (GEO) database and NCBI Pubmed related to SARS-CoV-2 infection in different types of human respiratory tract cells (Calu-3, doi:10.3390/biology9090260; ACE2-A549,GSE154613; a lung organoid model driving by human pluripotent stem cells, GSE155241; Primary human bronchial epithelial cells, GSE150819) which carried out in human cells (ex vivo) or human cell lines (in vitro). Genes with fold change>1.3 or <0.77 and p-values <0.05 were considered to be statistically significant.

### Functional analysis of differential expressed genes

The Go terms enrichment, KEGG pathways and protein-protein interaction analysis of the DEGs were performed using OmicsBean (http://www.omicsbean.cn) and Cytoscape v. 3.1.0 software (http://www.cytoscape.org/) [23] as a complementary and more comprehensive approach for identifying the central hub genes.

## RESULTS

### Overlapping and functional enrichment analysis of differential expression genes among different lung cells infected with SARS-CoV-2

To highlight the changes in cellular host genes in response to SARS-CoV-2 infection and further explore the mechanism of viral infection and pathogenesis, we searched the electronic databases (GEO databases or Pubmed) to identify the related studies. We included 4 studies in our qualitative study, they are all about transcriptome profiling of human respiratory tract cells (Calu-3, doi:10.3390/biology9090260; ACE2-A549, GSE154613; Lung organoid model driving by human pluripotent stem cells, GSE155241; Primary human bronchial epithelial cells, GSE150819).

Using Venn diagram analysis for the groups’ data, the following was found: there were 36 overlapping up-regulated and 0 overlapping down-regulated mRNAs among the four infection groups. (Figure 1A,B). We further performed a functional enrichment analysis using the 36 all up-regulated genes, and results revealed 2,506, 164, 226, and 102 terms associated with biological processes (BPs), cell components (CCs), molecular functions (MFs), and Kyoto Encyclopedia of Genes and Genomes pathways (KEGG), in which, 1,907, 78, 98, 39 terms was significantly enriched, respectively (Figure 2 A), top 10 enriched terms of BPs, CCs, MFs were presented in detail (Figure 2 B).

**Figure 1.**
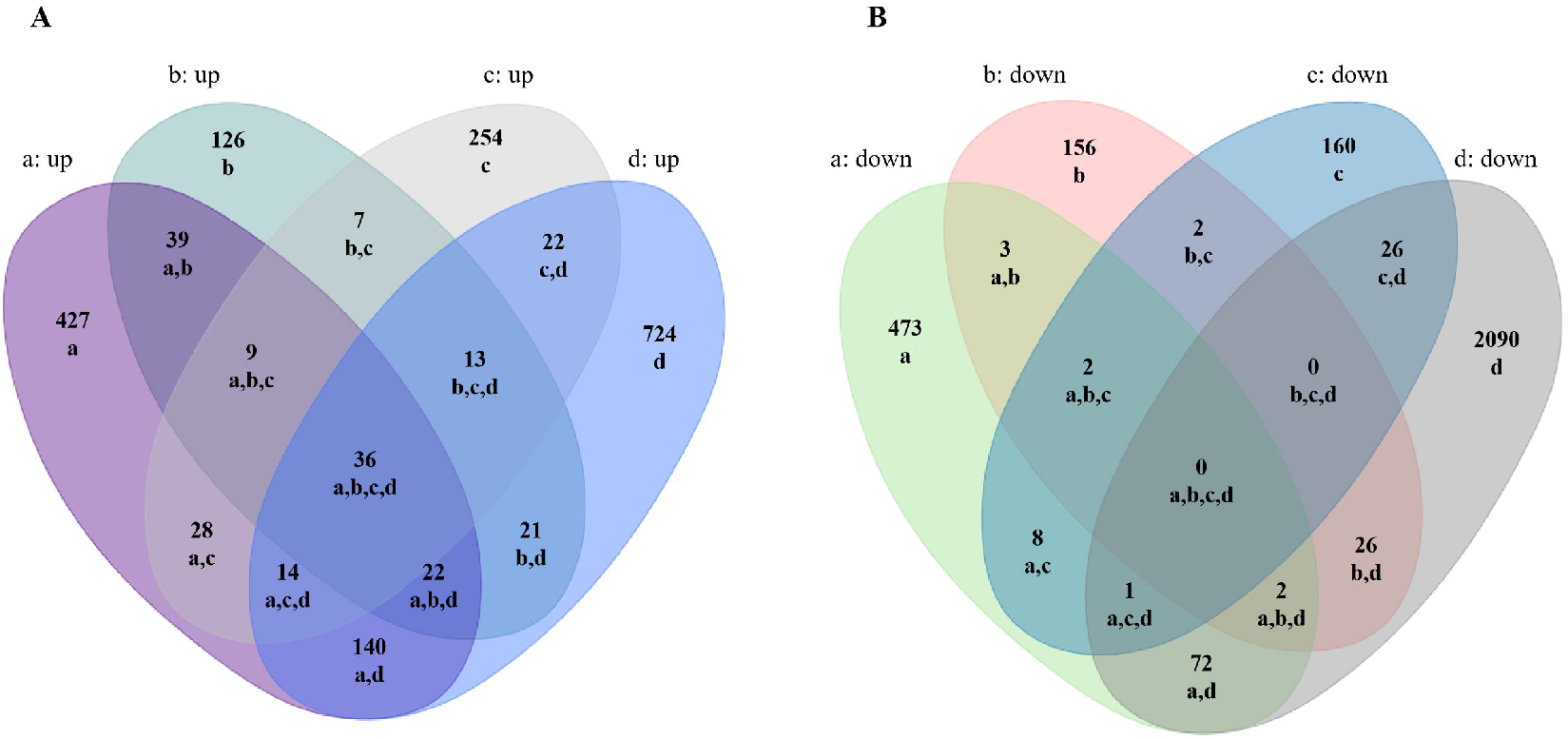
Comparative analysis of differentially expressed mRNAs in SARS-CoV-2 infected lung cells. A,B Venn analysis of all up/down expressed mRNAs among the four SARS-CoV-2 infection groups. a: Calu-3 infected with SARS-CoV-2 (doi:10.3390/biology9090260), b: ACE2-A549 infected with SARS-CoV-2 (GSE154613), c: Lung organoid model using human pluripotent stem cells infected with SARS-CoV-2 (GSE155241), d: Primary human bronchial epithelial cells infected with SARS-CoV-2 (GSE150819).

**Figure 2.**
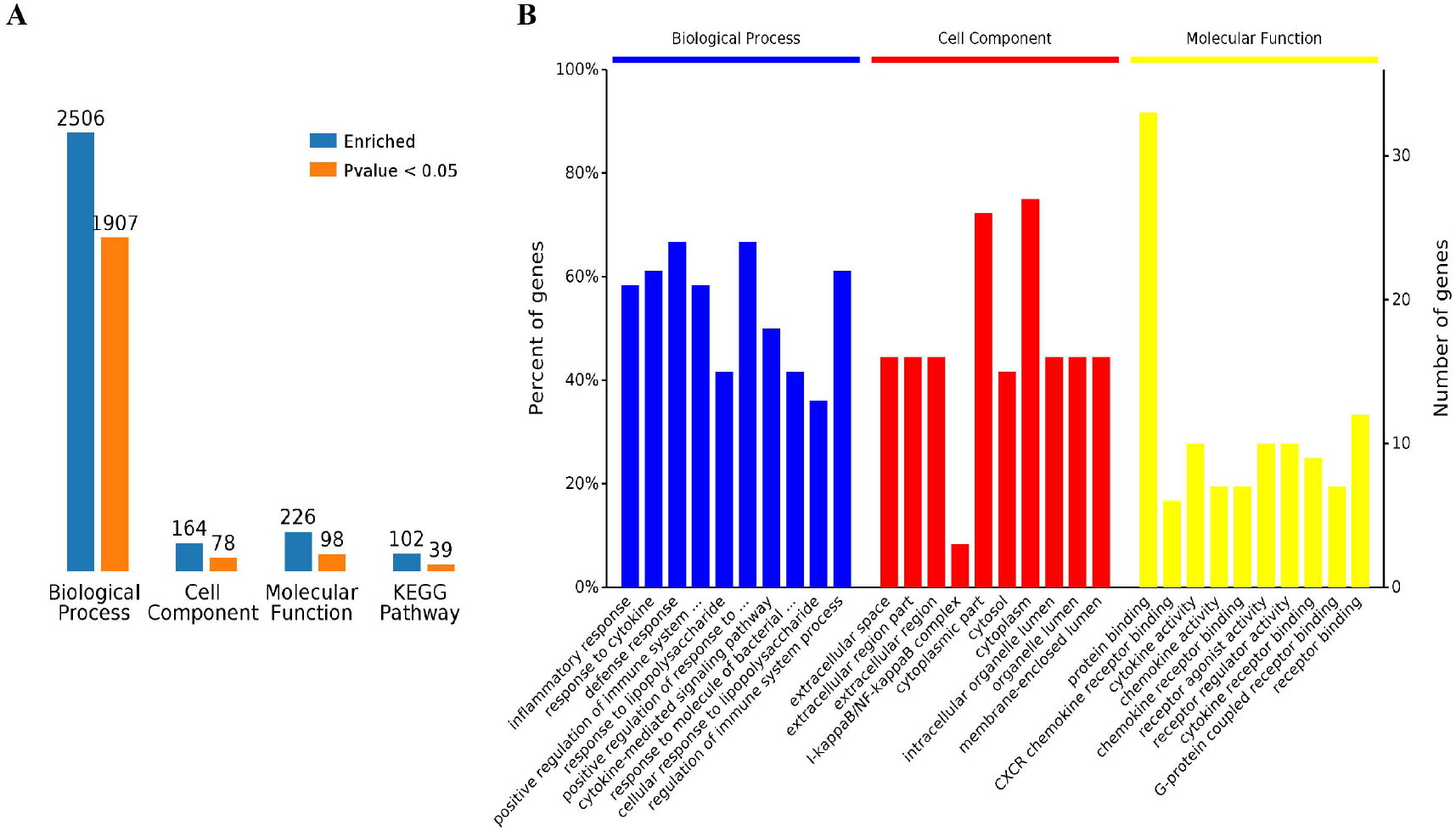
GO term enrichment analysis of SARS-CoV-2 induced up-regulated genes in all four groups. A, Number of enriched GO terms and KEGG pathways. B, Overview of significantly enriched biological processes, cell components, and molecular functions.

### Biological process analysis of Overlapping all up-regulated genes

The 1,907 different biological process GO terms were further analyzed using OmicsBean software. According to the different levels of the GO terms (Figure 3A, C), most differential genes were related to positive regulation of immune system process, regulation of immune system process and positive regulation of response to stimulus (level 3 GO), and continued subdivision of the grades showed that the differential genes were involved in the defense response, response to cytokine, cytokine-mediated signaling pathways (level 3-7 GO). According to the number of enriched genes (Figure 3B), they mainly enriched in signal transduction (29 differential genes), defense response (24 differential genes), regulation of cell communication (24 differential genes), response to organic substance (23 differential genes), multicellular organism development (22 differential genes) and cellular response to chemical stimulus (20 differential genes). A sorting of the different GO terms based on the P-value and gene number (Figure 3 C,D,E) revealed that these up-regulated genes are most likely involved in biological processes related to inflammatory response and response to cytokine, which indicates that SARS-CoV-2 infection could cause a strong host immune response.

**Figure 3.**
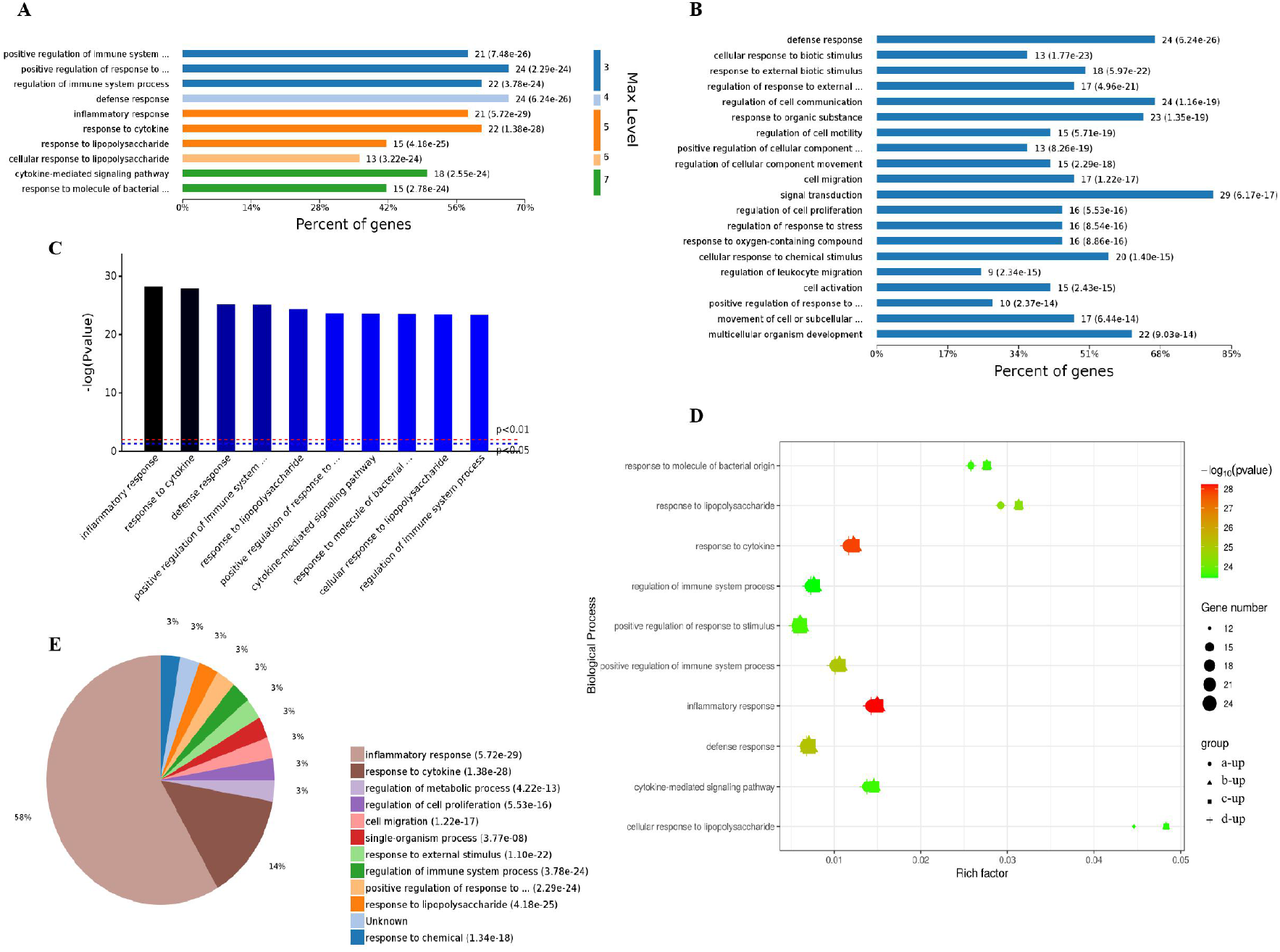
Analysis of 1907 biological process significantly enriched by all-up-regulated genes. A, Levels of enriched biological processes. B, Percent of genes associated with enriched biological processes. C, Top 10 of significantly enriched biological processes. D, Expressed genes associated with enriched biological processes. E, Pie chart of expressed proteins of enriched biological processes.

### Cell component analysis of all Overlapping up-regulated genes

The analysis of the 78 different cell components-related GO terms showed that according to the classification of the different GO levels (Figure 4 A), most up-regulated genes were related to continued subdivision identified extracellular region part, membrane-enclosed lumen (level 2 GO) and organelle lumen (level 3 GO), cytoplasm, cytoplasmic part and cytosol (level 4-6 GO). According to the number of enriched genes (Figures 4 B), the most abundantly enriched up-regulated genes were associated with cytoplasm (27 genes), intracellular organelle (22 genes), intracellular organelle part (20 genes) and vesicle (9 genes). By distinguishing the difference in the P-value of the GO terms and gene number (Figure 4 C,D), we found that these up-regulated genes were enriched in extracellular space and I-kappaB/NF-kappaB complex of cell components.

**Figure 4.**
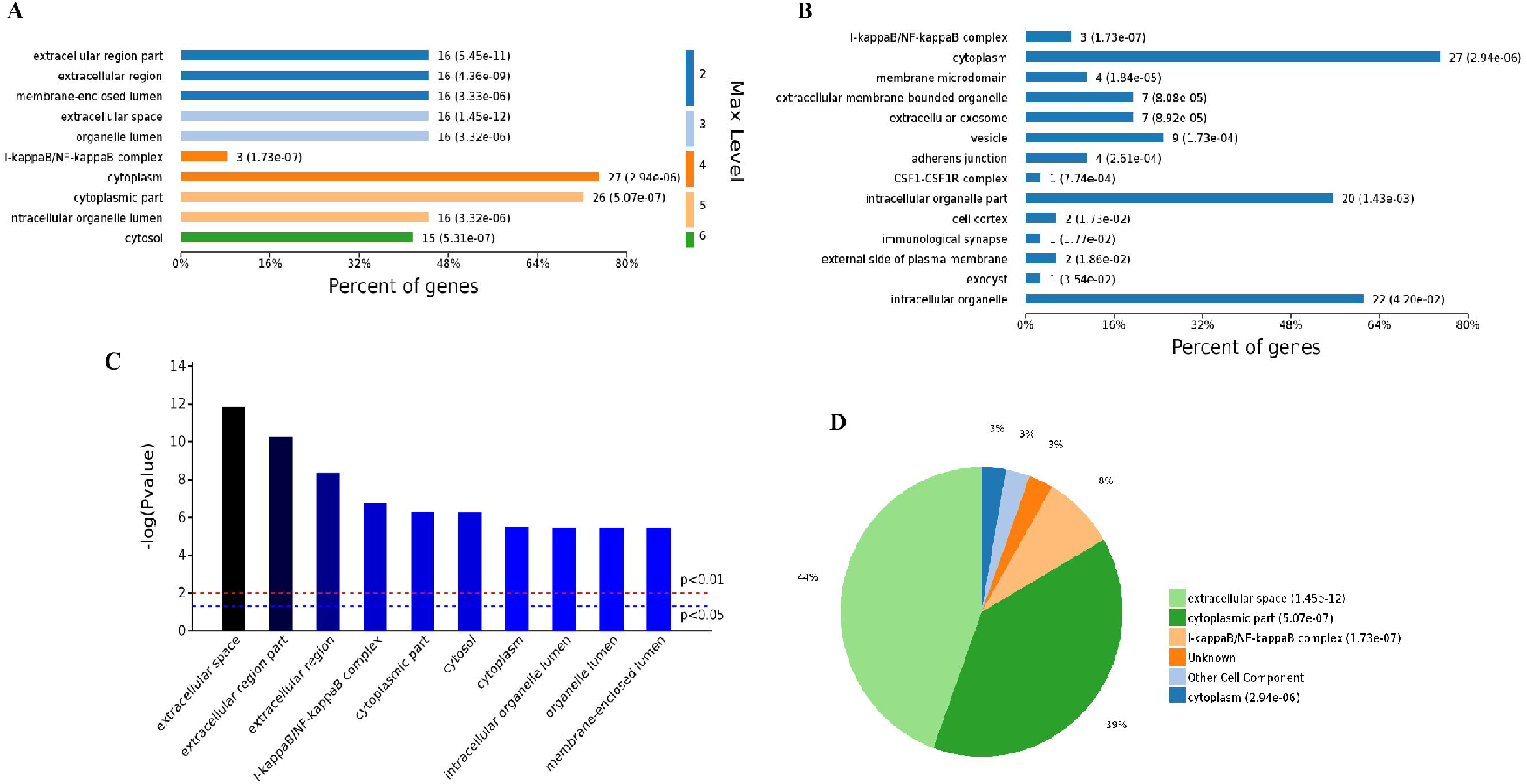
Analysis of 78 cell components significantly enriched by all up-regulated genes. A, Levels distribution of enriched cell components. B, Percent of genes associated with enriched cell components. C, Top 10 of significantly enriched cell components. D, Pie chart of expressed proteins of enriched cell components.

### Molecular function analysis of Overlapping all upregulated genes

The analysis of the 98 different molecular functions according to the number of the different GO classification levels (Figure 5A), the genes showing the highest differential expression are associated with protein binding (level 3 GO), and further subdivisions showed that the up-regulated genes are associated with receptor binding, cytokine receptor binding, G-protein coupled receptor binding, chemokine receptor binding and CXCR chemokine receptor binding (level 4-7 of GOs). In terms of the number of enriched genes (Figures 5 B) revealed that the most abundant up-regulated genes were enriched in receptor binding (12 genes), identical protein binding (10 genes) and enzyme binding (6 genes). Sorting the different GO terms based on ascending P-values and gene number (Figure 5 C,D) revealed that the genes showing the greatest differential expression are likely enriched in the molecular functions of the protein binding and cytokine activity.

**Figure 5.**
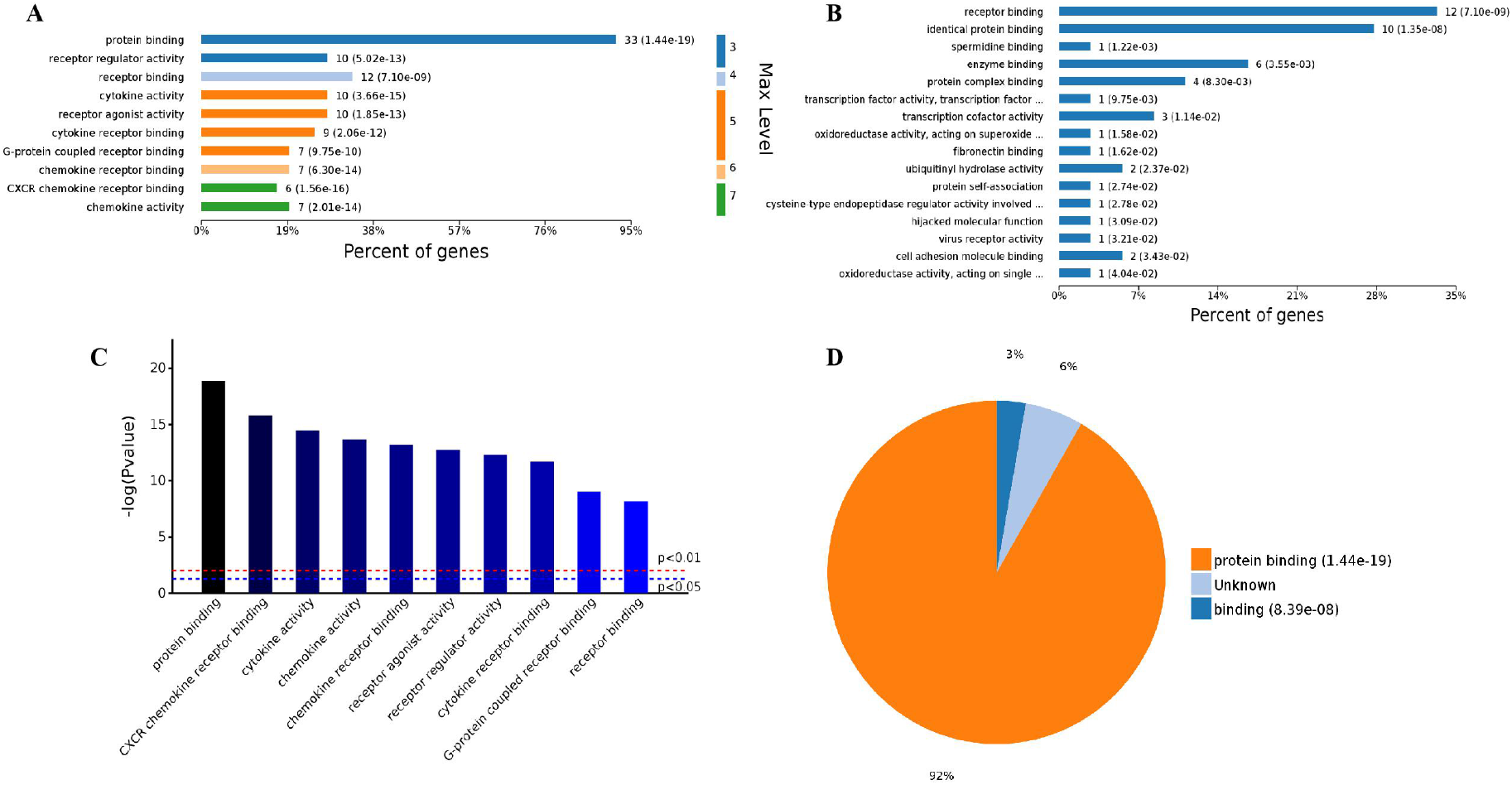
Analysis of 98 molecular function enriched by all up-regulated genes. A, Levels distribution of enriched molecular functions. B, Percent of genes associated with enriched molecular functions. C, Top 10 of significantly enriched molecular functions. D, Pie chart of expressed proteins of enriched molecular functions.

Based on the above results, it can be hypothesized that SARS-CoV-2 infection could activate the immune response in human respiratory tract cells. A statistical analysis of immune regulation GO terms and pathways (Supplemental Table 1-4) showed that many up-regulated genes are involved in.

### KEGG Pathway and protein-protein interaction Analysis of Overlapping all up-regulated genes

Through a KEGG pathway cluster analysis, we further found that the up-regulated genes mainly regulated the TNF signaling pathway, cytokine-cytokine receptor interaction, Viral protein interaction with cytokine and cytokine receptor and IL17 signaling pathway (Figure 6 A). Sorting the different KEGG pathways based on ascending P-values and gene number (Figure 6 B,C) revealed that the up-regulated genes mainly enriched in TNF signaling pathway, NF-kappa B signaling pathway and IL17 signaling pathway.

**Figure 6.**
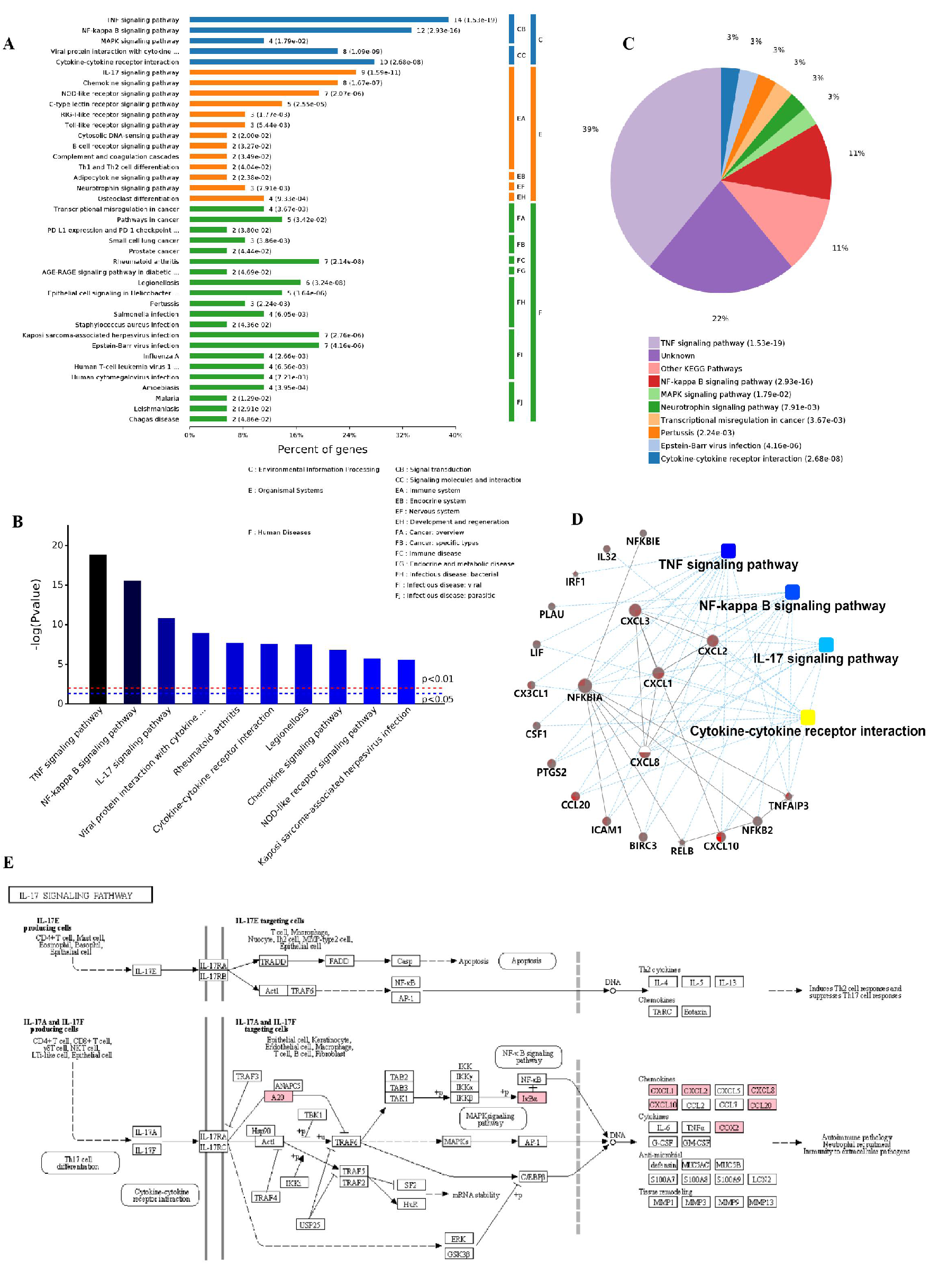
KEGG pathway cluster analysis and correlation analysis of all up-regulated genes. A, Percent of genes associated with enriched KEGG pathway. B, Distribution of enriched KEGG pathways. C, Pie chart of significant enriched KEGG pathway. D, Correlation analysis of up-regulated genes associated with key pathways. E, SARS-CoV-2 infection regulated the IL17 signaling pathway. The key up-regulated factors are marked in red.

A protein-protein interaction analysis (Figure 6 D) were also explored. The results showed that the key genes regulating these processes are CXCL1, CXCL2, CXCL3, CXCL8, CXCL10, IL32, CX3CL1, CCL20, IRF1, NFKB2 and NFKB1A, NFKB1E and their certain interaction relationship was also shown. Furthermore, an analysis of the most significant IL17 signaling pathway in KEGG (Figure 6 E) revealed that the TNFAIP3, NFKBIA (upstream) gene in the pathway and the CXCL1, CXCL2, CXCL8. CXCL10, CCL20 and PTGS2, (downstream) genes in the pathway, play an important role in immunity of host cells to SARS-CoV-2 infection. In addition, the details of NF-KAPPA B and TNF signaling pathway activated by SARS-CoV-2 infection were also shown in Figure S1-2. This finding is consistent with the results of the GO analysis, which showed that SARS-CoV-2 infection mainly activated the ability of respiratory tract cells to respond to the invasion of extracellular pathogens.

## DISCUSSION

As of 13 Oct 2020, a total of 37,423,660 confirmed cases of COVID-19 with 1,074,817 deaths had been reported [24], the case numbers are still increasing rapidly due to the second wave of epidemics worldwide. The underlying biological processes and mechanisms impacted by this viral infection of the host is still not clear. Analyzing the gene expression profiles of host cells infected with SARS-CoV-2 will be necessary to decipher the subcellular functions perturbed by this virus and to inform drug development strategies.

RNA-Seq has contributed to increase the knowledge relative to the biological and cellular processes involved during viral infection, especially in the context of an emergent pathogen [16,17], such as SARS-CoV-2. Venn analysis of several RNA-Seq assays increases the statistical power of samples and we applied this method for re-examining past evidence to better understand the pathogenesis of infections. In our current study, we explored recent SARS-CoV-2 infected cell-derived transcriptome datasets to dissect the cellular and molecular changes in response to SARS-CoV-2 infection. We finally included four available studies and analyzed the gene expression patterns using Venn analysis (Figure 1).

The RNA-Seq results of respiratory tract cells infected by SARS-CoV-2 showed that the up-regulated genes identified from the comparison of two groups such as CXCL1, CXCL2, CXCL3, CXCL8, CXCL10, IL32, CX3CL1, CCL20, IRF1, NFKB2 and NFKB1A, NFKB1E are involved in the regulation of TNF signaling pathway, NF-kappaB signaling pathway and IL17 signaling pathway. Tumor necrosis factor (TNF) was first described as a glycoprotein induced in response to endotoxin and has the capacity to kill tumor cells [25]. Then, TNF was shown to be a T helper type 1 (Th1) cytokine produced by several cell types, and to be involved in pro-inflammatory responses like interleukin (IL)-1β and IL-6 [26]. In concert with chemokines, TNF signaling pathway is involved in the regulation of inflammatory processes of infectious diseases including HIV (Human immunodeficiency virus), JEV (Japanese encephalitis virus), WNV (West Nile virus) and USUV (Ustuuvirus) infection [27-30]. The nuclear factor NF-kB pathway has also long been considered a prototypical pro-inflammatory signaling pathway, largely based on the role of NF-kB signaling pathway in the expression of pro-inflammatory genes including cytokines and chemokines [31]. Therefore, NF-κB has been considered as a target for new anti-inflammatory drugs for a long time. In our study, we found SARS-CoV-2 infection could activate NF-kB signaling pathway and up-regulate some pro-inflammatory genes, we further speculate drugs targeting the NF-kB signaling pathway may be one of the candidate drugs for the treatment severe COVID-19 patients.

It is worth mentioning that, IRF1 in respiratory tract cells was up-regulated by SARS-CoV-2 infection. IRF1 is a member of the IFN regulatory factor (IRF) family that plays an important role in immunomodulatory. IRF1 prominently participates in antiviral defense by regulating early expression of IFNs [32]. What’s more, as a transcription factor, IRF1 regulates constitutive expression of other antiviral genes by binding to their promoter directly. Studies revealed that IRF1 deficient mice are more susceptible to RNA viruses infections [33-37]. Thus, we think IRF1 may play a critical role in regulating constitutive antiviral gene networks to confer resistance against SARS-CoV-2 infections in human respiratory epithelial cells.

Inflammation is the first line of defense against tissue damage caused viral infection, involving activating both the innate and adaptive immune responses [38]. However, exuberant immune responses after viral infection, which also called cytokine storm, has been found to be associated with excessive levels of pro-inflammatory cytokines and also some tissue damage [39]. Cytokine storm has been proved to exist after several influenza viruses infection [40,41], as well as coronavirues [42-44], and contributes to acute lung injury or ARDS. Preliminary studies have revealed that SARS-CoV-2 infection could also trigger cytokine storm in severe cases, accompanied by an increase of various cytokines [45]. In the current study, we found that several cytokine related genes were upregulated after SARS-CoV-2 infection which could provide explanation about the cytokine storms associated with severe COVID-19 cases.

## CONCLUSIONS

Our results highlighted activation of TNF, NF-kappa B, Cytokine-cytokine interaction, IL17 signaling pathway and other inflammatory response as the main hallmark associated with SARS-CoV-2 infection in different types of human respiratory epithelial cells in vitro, further identified several differential expressed genes during the course of infection such as IRF1, CXCL1-3, IL32, CXCL10, CX3CL1, CCL20, NFKB2 and so on, which may serve as COVID-19 biomarkers. Interferon and inflammatory response to SARS-CoV-2 infection might provide explanation to cytokine storms associated with severe COVID-19 cases. However, precise mechanism in the host response to SARS-CoV-2 remains to be further investigated.

## DATA AVAILABILITY

We searched for RNA-Seq experiments deposited in the Gene Expression Omnibus (GEO) database and NCBI Pubmed related to SARS-CoV-2 infection in different types of human respiratory tract cells (Calu-3, doi:10.3390/biology9090260; ACE2-A549,GSE154613; a lung organoid model driving by human pluripotent stem cells, GSE155241; Primary human bronchial epithelial cells, GSE150819) which carried out in human cells (ex vivo) or human cell lines (in vitro).

## ACKNOWLEDGEMENTS

This work was supported by Shenzhen Natural Science Foundation (JCYJ20190809152415652), China Postdoctoral Science Foundation (2019M660836), the National Science and Technology Major Project (2018ZX10711001, 2017ZX10103011, 2018ZX09711003, and 2020YFC0841700), Shenzhen Science and Technology Research and Development Project (202002073000001).

## AUTHOR CONTRIBUTIONS

Yingxia Liu, Yu Wang, Minghui Yang and Luping Lei contributed to the study design. Minghui Yang, Luping Lei, Yang Yang and Jun Wang contributed to the manuscript and figures. Qiumei Cao, Xiao Jiang, Jinzhi Lai, Ling Qing and Kun Huang contributed to the proof reading. All the authors have read and approved the manuscript.

## SUPPLEMENTAL FILES

**Figure S1. The details of NF-KAPPA B signaling pathway activated by SARS-CoV-2 infection.** The key up-regulated factors are marked in red.

**Figure S2. The details of TNF signaling pathway activated by SARS-CoV-2 infection.** The key up-regulated factors are marked in red.

**Table S1. Summary of enriched Biological Processes.**

**Table S2. Summary of enriched Cell Components.**

**Table S3. Summary of enriched Molecular Functions.**

**Table S4. Summary of enriched KEGG pathways.**

