## Supplementary figures and images for "Transcriptome Profiling of different types of human respiratory tract cells infected by SARS-CoV-2 Highlight an unique Role for Inflammatory and Interferon Response"

### Figure S1.tif

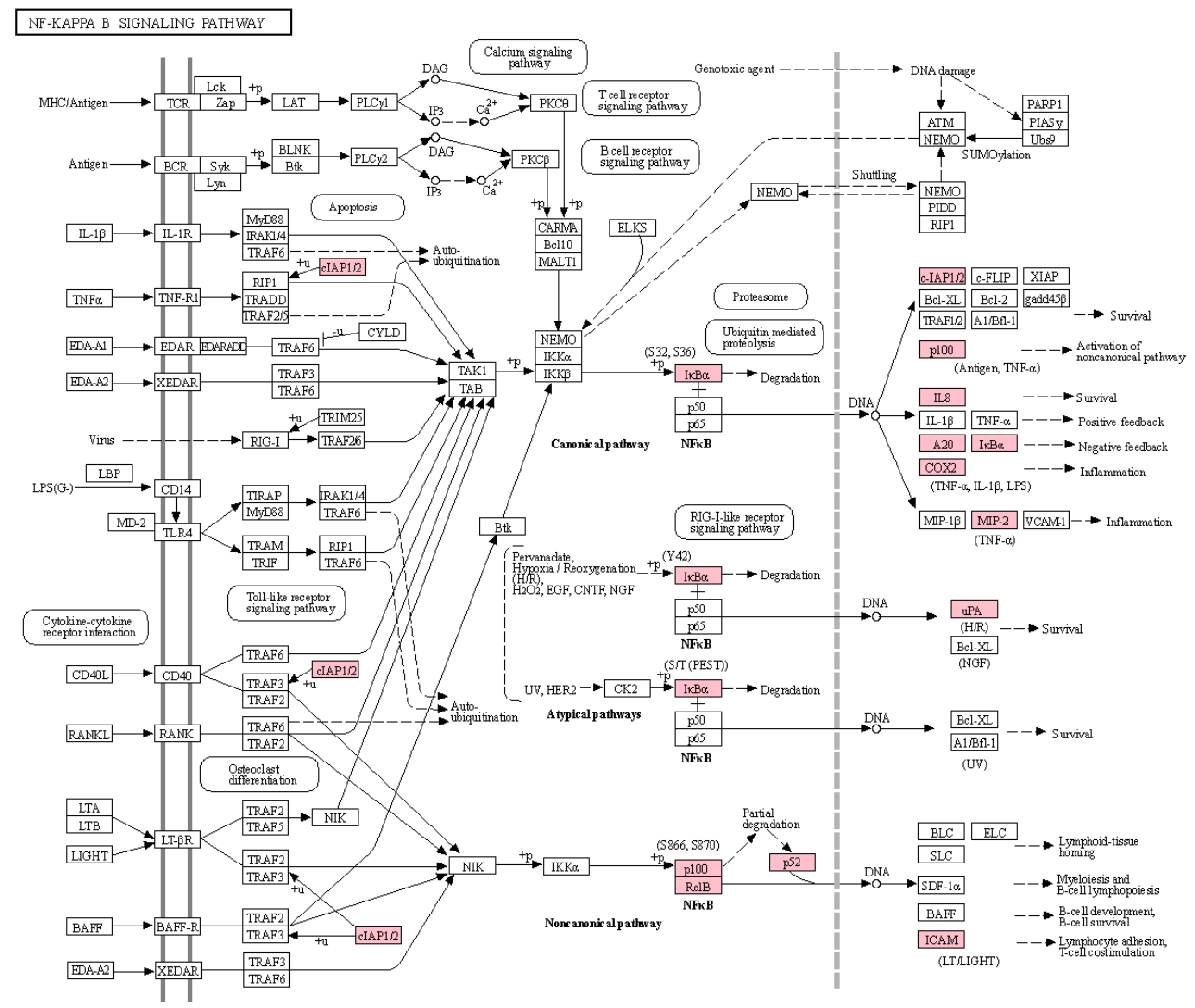

### Figure S2.tif

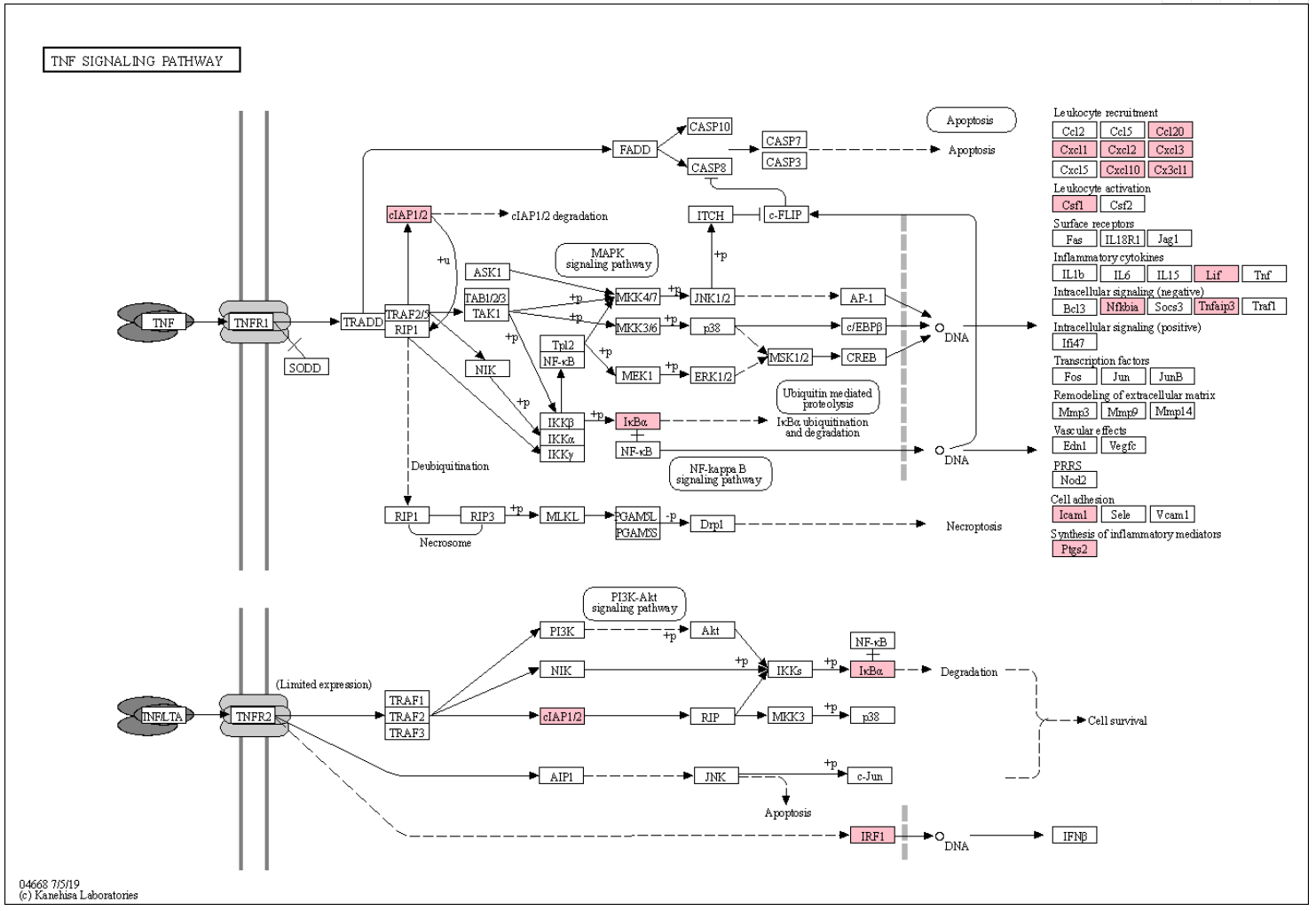
